# Enhanced Annotation of CD45RA to Distinguish T cell Subsets in Single Cell RNA-seq via Machine Learning

**DOI:** 10.1101/2023.05.23.541821

**Authors:** Ran Ran, Douglas K. Brubaker

## Abstract

T cell heterogeneity presents a challenge for accurate cell identification, understanding their inherent plasticity, and characterizing their critical role in adaptive immunity. Immunologists have traditionally employed techniques such as flow cytometry to identify T cell subtypes based on a well-established set of surface protein markers. With the advent of single-cell RNA sequencing (scRNA-seq), researchers can now investigate the gene expression profiles of these surface proteins at the single-cell level. The insights gleaned from these profiles offer valuable clues and a deeper understanding of cell identity. However, CD45RA, the isoform of CD45 which distinguish between naïve/central memory T cells and effector memory/effector memory cells re-expressing CD45RA T cells, cannot be well profiled by scRNA-seq due to the difficulty in mapping short reads to genes. In order to facilitate cell type annotation in T cell scRNA-seq analysis, we employed machine learning and trained a CD45RA^+/-^ classifier on single-cell mRNA count data annotated with known CD45RA antibody levels provided by cellular indexing of transcriptomes and epitopes sequencing (CITE-seq) data. Among all algorithms we tested, the trained support vector machine (SVM) with a radial basis function (RBF) kernel with optimized hyperparameters achieved a 99.96% accuracy on an unseen dataset. The multilayer Perceptron (MLP) classifier, the second most predictive method overall, also achieved a decent accuracy of 99.74%. Our simple yet robust machine learning approach provides a valid inference on the CD45RA level, assisting the cell identity annotation and further exploring the heterogeneity within human T cells.

## 1 Introduction

T cells play a pivotal role in adaptive immunity, serving as the cornerstone of the body’s defense mechanism against pathogens[1]. Each T cell clone has unique, sophisticatedly rearranged T cell receptors (TCRs) expressed on their surface, allowing them to bind to a specific group of antigens sourced from abnormal cells or foreign organisms and initiate an immune response[2]. Also, T cells are a remarkably heterogeneous population that can be further divided into subsets, such as CD4/CD8 T cells if by their surface glycoproteins, *αβ*/*γδ* T cell by their TCR chains, or naive/stem/memory/effector by their functions[3, 4, 5, 6]. Those subsets play different roles in maintaining immune system homeostasis and orchestrating immune response. Such heterogeneity underscores their versatility but also poses significant challenges to accurate cell characterization, which is important for developing strategies for ameliorating or reversing diseases.

Historically, immunologists have relied on protein-targeted techniques such as immunofluorescence, western blot, and flow cytometry to distinguish T cell subtypes in blood/peripheral tissue samples[1]. Those tools enable the characterization of cells based on a well-established set of surface protein markers, including but not limited to CD4, CD8, CD25, CD45, CD127, CCR7, CD62L, CD45RO, and CD45RA [3][7]. By analyzing the expression patterns of these markers, scientists have been able to classify T cells into helper T cells, cytotoxic T cells, regulatory T cells, memory T cells, etc.[8][9], and each is proved to have distinct functions and roles within the immune system. The development of techniques like single-cell RNA sequencing (scRNA-seq) has enabled the discovery of novel T-cell populations at an even higher resolution and provided a deeper understanding of their distinct functions in the immune system[10]. Specifically, scRNA-seq reports the counts of genes as the indicator of the transcription activities of that gene in every single cell in the sample[11]. With such a whole view of the cell features, under the assumption that the same type of cells has a similar pattern of gene expression, unsupervised clustering on the feature matrix should be able to generate clusters that are the union of cells of presumably the same cell type, which provides insights into cell subtypes identification[12].

Despite these advances, certain limitations persist in the application of the scRNA-seq for T-cell studies. One such challenge is the accurate profiling of CD45RA, an isoform of the CD45 protein. CD45RA is the result of alternative splicing of CD45 and serves as a canonical marker to distinguish between naïve T/central memory T (TCM) cells (naive T is CD45RA^+^ while TCM is CD45RA^-^) and effector memory T (TEM)/effector memory re-expressing CD45RA T (TEMRA) cells[13]. However, the expression of CD45RA mRNA can be challenging to report using scRNA-seq techniques because most of them rely on next-generation sequencing (NGS) platforms that fragment the cDNA of mRNA and obtain gene information by mapping the short reads to the human genome in the downstream analysis[14][15]. Therefore, it is not uncommon for CD45RA information to be missing or less reliable in the output count matrix from scRNA-seq, which adds difficulties to the optimal annotation of T cell subsets identities. To overcome this limitation, researchers have turned to alternative approaches, including the use of multi-omic techniques that combine scRNA-seq with other technologies, such as cellular indexing of transcriptomes and epitopes by sequencing (CITE-seq)[16]. CITE-seq enables simultaneous measurement of surface protein expression and gene expression at the single-cell level, offering a more comprehensive view of cellular identity. Instead of reporting the level of CD45RA mRNA, CITE-seq can measure the product, alias the CD45RA protein, along with the RNA profiles.

Still, a method to better infer the CD45RA level in conventional scRNA-seq experiments is needed for studying human immunology. Retrospectively, there is likely important information about CD45RA identified in studies performed prior to the invention of CITE-seq, where such information would re-contextualize and enhance the interpretation of previously collected data. Prospectively, even as CITE-seq gains popularity, there will likely be biological use cases where the technique could be more feasible and cost prohibitive. Such methods for inferring CD45RA status are potentially important for immunology in the future and may need to be expanded to other markers besides CD45RA. Given the production of protein is the result of a series of highly-orchestrated gene expressions, an intuitive hypothesis is that CD45RA^+^ cells should have a different pattern in mRNA counts compared to CD45RA^-^ cells. Therefore, we employed machine learning approaches and trained multiple CD45RA^+/-^ classifiers on the NGS single-cell mRNA count matrix with known CD45RA antibody levels reported by CITE-seq to facilitate cell type annotation in T cell NGS scRNA-seq analysis.

## 2 Method

### 2.1 Data Acquisition

Raw CITE-seq counts of 5559 healthy adult peripheral blood mononuclear cells labeled with CD45RA antibody level were obtained from NCBI Gene Expression Omnibus (GEO) series GSE144434 as the training/testing dataset[17]. To test the performance and robustness of models on NGS scRNA-seq data generated by different experiments, the Cytometry by Time of Flight (CyTOF)-sorted scRNA-seq raw counts of CD45RA^-^ T cells in healthy human blood from study GSE150132[18].

### 2.2 Single Cell RNA-seq Data Processing

The downstream analysis was done with scanpy (v1.9.1)[19] on Python and Seurat (v4.2.0) on R[20]. Cells with CD19 or without CD3D/E/G/Z expression were discarded only to keep the profiles of T cells. Low-quality cells were already dropped by the data provider based on their extremely low UMI counts, high mitochondrial gene counts, and low number of uniquely expressed genes. In this study, cells with less than 5% of the ribosomal gene count were also dropped. Genes expressed in less than 2 cells in each dataset were excluded. Multiplets were removed by Scrublet (v0.2.3)[21]. After basic quality control, the 11,245 genes’ transcriptional profile in 4,000 cells was obtained. The cell cycle score was calculated based on the expression of cell cycle-related genes as previously described[22]. The dataset was normalized by SCTransform (v0.3.5) on R[23]. Cell mitochondrial fraction, and the difference between the S phase score and the G2M phase score, as proposed as the representation of the cell cycle score, was regressed during the normalization. 3000 highly variable features were re-calculated for the combined dataset and used to perform the principal component analysis (PCA, 50 pcs). Leiden clustering (resolution = 0.5) was performed based on the computed neighborhood graph of observations (UMAP, 50 pcs, size of neighborhood equals 15 cells) to reveal the general subtypes of the T cells[13][24]. Partition-based graph abstraction (PAGA) based on the Leiden clusters, with a threshold of 0.2, was used to initialize the uniform manifold approximation (UMAP, 50 pcs, min_dist = 0.01, spread = 2, n_components=2, alpha=1.0, gamma=1.0) to facilitate the convergence of manifold[25].

### 2.3 Determine CD45RA Positive/Negative Label

The CD45RA level reported by CITE-seq is derived from the counts of unique DNA-barcoded sequences associated with CD45RA antibodies[16]. It has been preprocessed by the data provider. The Otsu method, also known as Otsu’s thresholding, in the skimage library[26] is used to calculate the optimum threshold *t\** of CD45RA level to separate positive and negative cells. Specifically, the Otsu method aims to find a threshold *t\** that maximizes the between-class variance while implicitly minimizing the within-class variance. Since the overlapping observations (i.e., cells that had a CD45RA level close to *t\**) were hard to be assigned a positive/negative label, to reduce the false positivity/negativity in CD45RA^+/-^ labeling, CD45RA^+^ cells were defined as cells having a CD45RA level >= *t\** + 0.5, and CD45RA^-^ cells were defined as cells having a CD45RA level <= *t\** − 0.5. In other words, overlapping observations were not used in the model training.

### 2.4 Differentailly Expressed Genes Identification

Differential expression (DE) analysis in Monocle 3[27] involves fitting a generalized linear model (GLM) to the SCTransform-corrected counts. The quasi-Poisson distribution is used as the noise model to account for both the mean and variance in the data.

For a given gene, let *y*_*ij*_ represent the observed expression level for cell *i* in condition *j*, where *i* = 1, 2, …, *n*, and *j* = 1, 2 (two conditions, CD45RA^+/-^). The quasi-Poisson GLM can be written as:

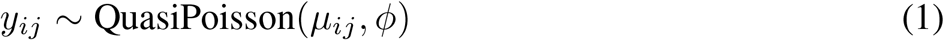

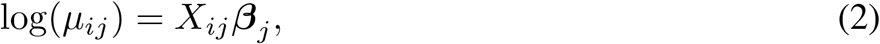

where *μ*_*ij*_ is the expected expression level for cell *i* in condition *j, ϕ* is the dispersion parameter, *X*_*ij*_ is the design matrix representing covariates (e.g., experimental conditions, batch effects), and ***β***_*j*_ is the vector of regression coefficients for condition *j*.

The link function is the natural logarithm, which maps the expected expression level (*μ*_*ij*_) to the linear predictor (*X*_*ij*_***β***_*j*_), helping ensure that the expected expression level is always non-negative.

The dispersion parameter (*ϕ*) accounts for the overdispersion in the data, which occurs when the variance is greater than the mean. In the quasi-Poisson model, the variance is modeled as:

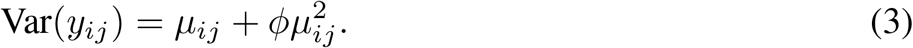

For each gene, the GLM is fit to the data using maximum likelihood estimation, which involves finding the ***β***_*j*_ and *ϕ* values that maximize the likelihood of the observed data.

A likelihood ratio test is performed to test for differential expression between the two conditions. This compares the likelihood of the data under the full model (with separate ***β***_*j*_ values for each condition) to the likelihood under the null model (with the same ***β***_*j*_ value for both conditions). The test statistic is calculated as follows:

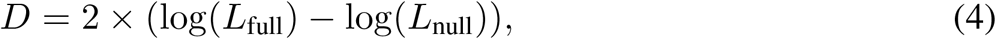

where *D* follows a chi-square distribution with degrees of freedom equal to the difference in the number of parameters between the full and null models. A q-value is calculated from the test statistic, and genes with q-values below 0.05 are considered DE genes.

### 2.5 Feature Selection

After splitting data into training and testing sets, features were selected from two sources. The first source was the DE genes between CD45RA^+/-^ groups. The fold change of a DE gene’s level in one group with respect to the other could be quantified to describe the extent of differential expression. DE genes with a fold change above an arbitrary threshold, *t*_DE_, were selected as input features for the classifier. By adjusting *t*_DE_, it became possible to fine-tune the input features’ degree of distinction across CD45RA^+/-^.

Mathematically, given the SCTransform-corrected gene expression values in CD45RA^+^ (*C*_1_) and CD45RA^-^ (*C*_2_), the fold change (*FC*) for a specific gene can be calculated as follows:

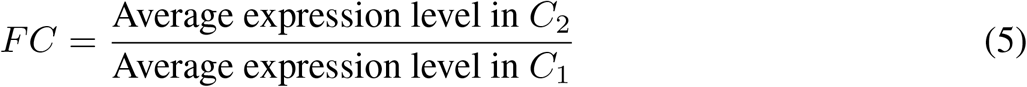

The log2-transformed fold change is defined as:

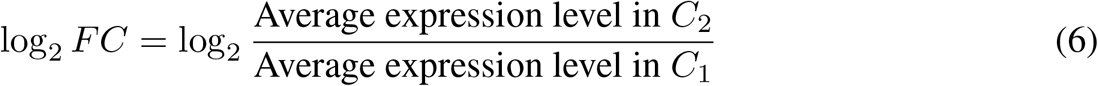

In the context of the quasi-Poisson GLM, the log2 fold change can be estimated by calculating the difference in the estimated coefficients, ***β***_*j*_, between the two conditions:

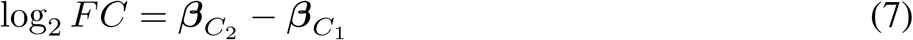

The second source for feature selection comes from previous biological knowledge. Heterogeneous nuclear ribonucleoprotein L-like (hnRNPLL) is an RNA-binding protein that plays a crucial role in alternative splicing, a process where pre-messenger RNA (pre-mRNA) is rearranged to produce different mRNA molecules and, consequently, various protein isoforms. hnRNPLL has been implicated in the regulation of CD45 alternative splicing, specifically in the generation of the CD45RA isoform. hnRNPLL binds to specific RNA sequences in the CD45 pre-mRNA, promoting the inclusion or exclusion of specific exons. The binding of hnRNPLL to CD45 pre-mRNA has been shown to enhance the exclusion of variable exons, leading to the generation of the CD45RA isoform[28]. In this manner, hnRNPLL helps regulate the expression of CD45 isoforms and contributes to the proper functioning of the immune system.

Therefore, we hypothesized that hnRNPLL and genes correlated with it may be indicative of the CD45RA status. Spearman correlation coefficients of every gene’s expression with hnRNPLL’s expression were calculated. Genes that had an absolute correlation coefficient with hnRNPLL in the *P*_hn_ percentile with a p-value < 0.05 were selected as input features. Similar to the DE genes selection, the extent of the input features correlate with hnRNPLL can also be optimized later by adjusting *P*_hn_.

### 2.6 Classifier Training

#### 2.6.1 Support Vector Machine (SVM)

SVM is a supervised learning method often used for classification and regression problems[29]. It aims to find the optimal hyperplane to classify the data points while maximizing the margin, the closest distance from data points from each class to the hyperplane. SVM can efficiently process sparse data in high-dimensional spaces by relying on the inner product between feature vectors. Such a technique known as the “kernel trick” allows SVM to implicitly map data into higher-dimensional feature spaces for more effective linear decision boundaries. Furthermore, maximal margin separation enables SVM to find the optimal hyperplane in high-dimensional spaces, which helps generalize well to unseen data. Lastly, SVM incorporates regularization to prevent overfitting.

Training data was fitted to an SVM with a Radial Basis Function (RBF) kernel and another SVM with a linear kernel using scikit-learn (v1.1.2) in Python[30]. For the SVM with RBF kernel, *t*_DE_ and *P*_hn_ that threshold the input features, along with the cost *C*_SVM_ and the RBF kernel coefficient *γ* were optimized by a Bayesian Optimizer using Bayesian Optimization in Python[31], aiming to achieve the optimal 5-folds cross-validation averaged training accuracy in 30 iterations. *t*_DE_ ∈ (1, 4) or (−4, −1), *P*_hn_ ∈ (1*e* − 6, 0.1), *C*_SVM_ ∈ (1*e* − 6, 100), *γ* ∈ (1*e* − 6, 2). For the linear kernel SVM, no kernel coefficient was needed.

Eventually, the final SVM was trained on the feature-optimized training set and took the optimized hyperparameters. It was then used to predict the CD45RA label of testing data, unseen CD45RA^-^ data. The accuracy of the prediction on datasets with known CD45RA labels was reported, and the performance on classifying single-cell data of two SVMs were compared.

#### 2.6.2 Logistic Regression (LR)

Logistic regression is a classic and relatively simple binary classifier[32]. It can handle high-dimensional data since it relies on finding the optimal decision boundary, which is not affected by the “curse of dimensionality” in the same way as distance-based measures[33]. Also, LR is adept at handling the sparse scRNA-seq counts due to its ability to incorporate regularization, which encourages sparse solutions and helps in feature selection.

The training data was fitted to an LR classifier with a threshold for the predicted probabilities of 0.5 using scikit-learn (v1.1.2) in Python[30]. *t*_DE_ and *P*_hn_ that threshold the input features, along with the regularization strength C were optimized by a Bayesian Optimizer using Bayesian Optimization in Python[31], aiming to achieve the optimal 5-folds cross-validation averaged training accuracy in 30 iterations. *t*_DE_ ∈ (1, 4) or (−4, −1), *P*_hn_ ∈ (1*e* − 6, 0.1), *C*_LR_ ∈ (1*e* − 6, 2).

Eventually, the final LR classifier was trained on the feature-optimized training set and took the optimized hyperparameters. It was then used to predict the CD45RA label of testing data, unseen CD45RA^-^ data. The accuracy of the prediction on datasets with known CD45RA labels was reported.

#### 2.6.3 Support Vector Machine Stacked Logistic Regression

Stacking an LR model and an SVM together can potentially create a more powerful classifier by leveraging the strengths of both models[34]. LR is a linear model that works well when the decision boundary between classes is relatively linear. At the same time, SVM, particularly with non-linear kernels like RBF, can capture more complex decision boundaries.

We trained a meta LR using the predictions of the LR and SVM classifiers as input features and the true CD45RA^+/-^ labels as the target variable using scikit-learn (v1.1.2) in Python[30]. *t*_DE_ and *P*_hn_ that threshold the input features, along with the regularization strength *C*_LR_ from LR, the cost *C*_SVM_ and the RBF kernel coefficient *γ* from SVM were optimized by a Bayesian Optimizer using Bayesian Optimization in Python[31], aiming to achieve the optimal 5-folds cross-validation averaged training accuracy in 30 iterations. *t*_DE_ ∈ (1, 4) or (−4, −1), *P*_hn_ ∈ (1*e* − 6, 0.1), *C*_SVM_ ∈ (1*e* − 6, 100), *γ* ∈ (1*e* − 6, 2), *C*_LR_ ∈ (1*e* − 6, 2). Since the meta LR only had two features, which were the predictions by SVM and LR, its hyperparameter was not optimized.

The final model was trained on the feature-optimized training set and took the optimized hyperparameters. It was then used to predict the CD45RA label of testing data, unseen CD45RA^-^ data. The accuracy of the prediction on datasets with known CD45RA labels was reported.

#### 2.6.4 Multilayer Perceptron

The multilayer perceptron (MLP) is an artificial neural network characterized by its multiple layers of interconnected neurons[35]. This machine learning model has proven its mettle in various applications, including binary classification tasks[36, 37, 38]. In comparison to LR, the MLP excels in its ability to model non-linear relationships. While LR is effective in linear contexts, its simplicity restricts its capabilities in handling more complex data. The MLP, on the other hand, can learn non-linear patterns with ease, providing excellent performance in cases where linearity cannot be assumed. When it comes to SVM, while SVM can handle non-linear data through the use of kernels, it lacks the flexibility and scalability inherent to the MLP’s architecture. The MLP can be fine-tuned to a wide range of classification problems by adjusting the number of hidden layers and neurons. This adaptability makes the MLP more robust for tackling diverse and challenging datasets.

For simplicity, we used the optimized features in linear and RBF kernel SVM as the input features for two MLPs, respectively. The training data was fitted to an MLP threshold for the predicted probabilities of 0.5 using tensorflow (v2.12.0) in Python[39]. ReLu was used as the activation function for all layers except the output layer, which used sigmoid as its activation function. The epoch number was 15, and the batch size was 32 when building the MLP. The number of layers, the number of neurons, the learning rate, and the dropout rate were optimized by a Bayesian Optimizer with the aim of achieving minimal average binary cross-entropy of the 5-fold cross-validation in 30 iterations. *N*_layers_ ∈ (1, 3), *N*_nodes_ ∈ (16, 128), *η* ∈ (1*e* − 4, 0.01), *ρ* ∈ (0.1, 0.5). The final MLP was trained on the feature-optimized training set and took the optimized hyperparameters. It was then used to predict the CD45RA label of testing data, unseen CD45RA^-^ data. The accuracy of the prediction on datasets with known CD45RA labels was reported.

## 3 Results

### 3.1 Feature Interpretation

We found NPDC1, AQP3, CCR6, IL7R, DPP4, HES6, CD28, and another 116 genes were qualified as differentially expressed in CD45RA^+^ cells compared to CD45RA^-^ cells under the standard that is often used in general scRNA-seq studies (| log2FC | >= 2, q-value <= 0.05) (Figure 1A, Table S1). Among these DE genes, we found well-studied naïve/memory signatures like CD28 and IL7R[40]. Specifically, CD28 is a costimulatory receptor expressed on naive T cells, essential for T cell activation[41]. Interleukin 7 Receptor (IL7R, alias CD127) is a receptor that binds to interleukin-7 (IL-7) and plays a crucial role in the development and homeostasis of naive T cells[42]. Although previous literature described both naïve and (stem) central memory T cells express CD28 and IL7R, the difference in their GLM coefficients indicates levels of these genes in CD45RA^+/-^ cells differ.

**Figure 1.**
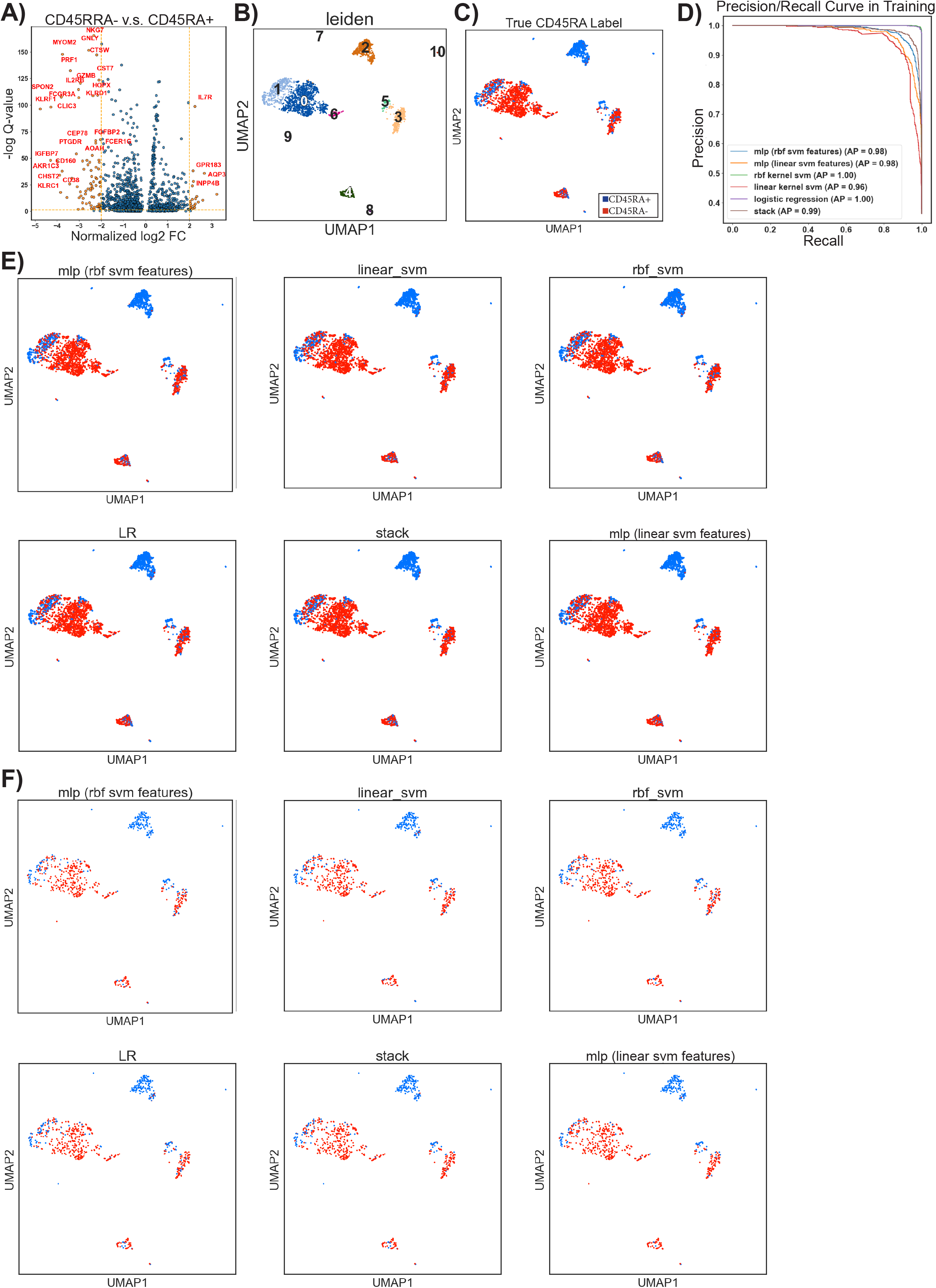
Overview of the features, the data, and the prediction made by classifiers. A) Volcano plot of the differentially expressed genes. x-axis: gene expression level log2 fold change in CD45RA-cells with respect to CD45RA+ cells. y-axis: -log10 q-value (false discovery rate) of gene’s fold change in CD45RA- with respect to CD45RA+. The threshold of the magnitude of the DE genes’ log2 FC across CD45RA+/- to be considered distinguished enough to be reported was set at 2. B) Uniform manifold approximation and projection (UMAP) of cell subsets in the CITE-seq data. Colors represent different Leiden clusters. C) Visualization of the CD45RA true label in the CITE-seq data. D) Precision/Recall curve during the training of all 6 classifiers. AP, Average Precision. E), F) Visualization of classifiers’ predictions in the E) training data and F) testing data embedded on the UMAP coordinates. The first subplot in each plot, as its title “True CD45RA Label” indicates, shows the true CD45RA label of the training/testing data.

Genes reported to be highly associated with T-cell differentiation, such as CD40LG, CTLA4, TBX21, and IL2RB, were also found in the DE genes. CD40LG encodes CD40 Ligand, a protein activating and regulating the immune system, including T cells[43]. CTLA4 translates into Cytotoxic T-Lymphocyte Associated Protein 4, a protein that functions as an immune checkpoint and negatively regulates T-cell activation[44]. TBX21, alias the gene of T-box 21, is a transcription factor involved in the differentiation of T cells[45]. It was expected to see DE analysis report these genes, given they are considered highly indicative of the activation of T cells and partially in coordination with the expression of CD45RA.

We also found Neural Proliferation Differentiation and Control 1 (NPDC1) and aquaporin 3 (AQP3) had the highest two log2 FC in CD45RA^-^ group compared to CD45RA^+^ group. NPDC1 is a protein-coding gene implicated in regulating cell proliferation, differentiation, and apoptosis in various cell types. Recently, it was reported to be a prognostic immune gene in a model that predicts the outcome of acute myeloid leukemia (AML)[46]. However, its specific role in T cell activation is not well-defined. AQP3 is a water channel protein that facilitates the transport of water and small solutes, such as glycerol, across cell membranes. AQP3 is expressed in various tissues, including the skin, kidneys, and gastrointestinal tract, and is involved in diverse physiological processes, such as water balance and skin hydration[47]. Its role in immune cells, specifically T cells, is not well-established, yet we did find its expression was highly overlapped with CD45RA^-^, suggesting CD45RA^-^ have a different metabolism compared to CD45RA^+^ cells.

When determining the second source for feature selection, hnRNPLL’s expression was found to correlate with the transcription of HNRNPA2B1, a member of the hnRNP family, which plays a role in pre-mRNA processing, along with other 1600 genes that had a p-value <= 0.05 (Table S2). Although HNRNPA2B1 has been reported to participate in alternative splicing[48], no direct evidence exists that it is associated with CD45 splicing specifically. Given its correlation with hnRNPLL, we suggest that it is possible that HNRNPA2B1 is also engaged in the CD45 splicing, inviting further investigation.

### 3.2 Performance of Classifiers on the Testing Set and Unseen Data

The thresholds used for feature selection and the resultant features of all 6 classifiers (Table 1, S3) are recorded. The final values of the optimized hyperparameters (Table 2) are also reported. The accuracy (Table 3), precision (Table 4), and recall (Table 5) in training, testing, and the unseen datasets were used as metrics to evaluate the performance of classifiers. We found that all those simple classifiers achieved a reasonably good accuracy (>85%) in training, testing, and unseen, and the SVM with an RBF kernel outperformed the other 4 classifiers in terms of all three types of accuracy with a relatively parsimonious selection of features.

**Table 1:**
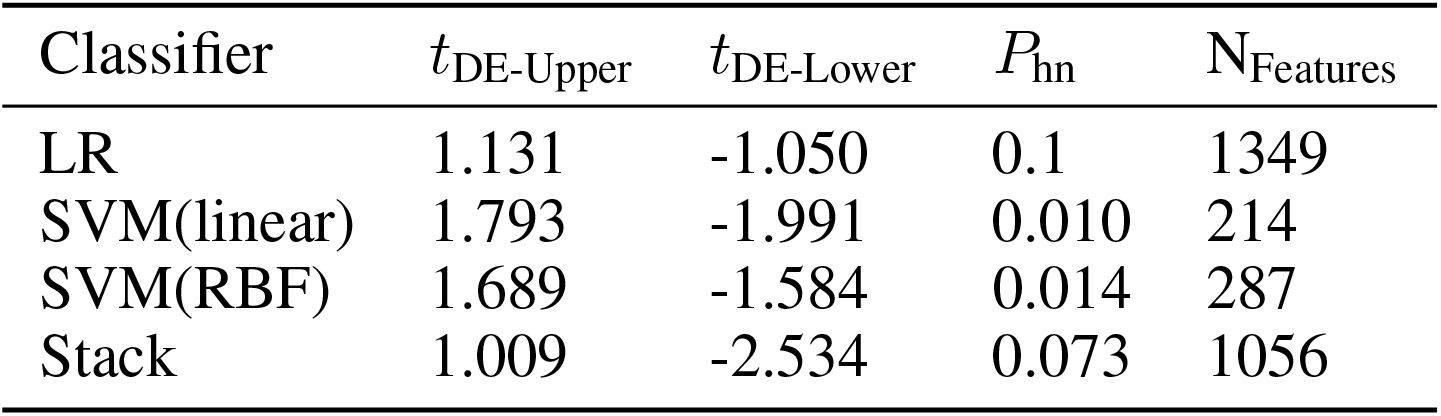
Feature Selection Parameters

| Classifier | $t_{DE-Upper}$ | $t_{DE-Lower}$ | $P_{hn}$ | $N_{Features}$ |
| --- | --- | --- | --- | --- |
| LR | 1.131 | -1.050 | 0.1 | 1349 |
| SVM(linear) | 1.793 | -1.991 | 0.010 | 214 |
| SVM(RBF) | 1.689 | -1.584 | 0.014 | 287 |
| Stack | 1.009 | -2.534 | 0.073 | 1056 |

**Table 2:** The Final Values of Hyperparameters

| Classifier | Hyperparameters |
| --- | --- |
| LR | $C_{LR} = 0.595$ |
| SVM(linear) | $C_{SVM} = 48.829$ |
| SVM(RBF) | $C_{SVM} = 54.343, \gamma = 0.069$ |
| Stack | $C_{SVM} = 1e-6, C_{LR} = 0.054$ |
| MLP (linear SVM features) | $N_{layers} = 1e-4, N_{nodes} = 0.1, \eta = 3, \rho = 105$ |
| MLP (RBF SVM features) | $N_{layers} = 2.262e-4, N_{nodes} = 0.1, \eta = 1, \rho = 87$ |

**Table 3:** Accuracy of Predictions in 3 Datasets

| Classifier | Training (%) | Testing(%) | Unseen(%) |
| --- | --- | --- | --- |
| LR | 99.31 | 88.78 | 88.62 |
| SVM(linear) | 92.89 | 86.59 | 99.60 |
| SVM(RBF) | 99.45* | 89.94* | 99.96* |
| Stack | 96.17 | 88.78 | 94.04 |
| MLP (linear SVM features) | 93.22 | 87.32 | 99.74 |
| MLP (RBF SVM features) | 95.48 | 89.21 | 97.63 |

**Table 4:** Precision of Predictions in Training and Testing Datasets

| Classifier | Training (%) | Testing(%) |
| --- | --- | --- |
| LR | 99.39 | 88.33 |
| SVM(linear) | 92.38 | 88.56 |
| SVM(RBF) | 99.60* | 90.20 |
| Stack | 97.04 | 90.87 |
| MLP (linear SVM features) | 94.70 | 92.31* |
| MLP (RBF SVM features) | 92.91 | 88.76 |

**Table 5:** Recall of Predictions in Training and Testing Datasets

| Classifier | Training (%) | Testing(%) |
| --- | --- | --- |
| LR | 98.69 | 82.85 |
| SVM(linear) | 87.65 | 76.28 |
| SVM(RBF) | 98.90* | 83.94* |
| Stack | 92.27 | 79.93 |
| MLP (linear SVM features) | 86.14 | 74.45 |
| MLP (RBF SVM features) | 94.78 | 83.58 |

Specifically, during the hyperparameter optimization, the LR classifier’s 5-fold cross-validation accuracy ranged from 62.77% to 90.42%, with an average of 86.79%. The linear kernel SVM’s 5-fold cross-validation accuracy ranged from 64.01% to 88.20%, with an average of 84.98%. The RBF kernel SVM’s 5-fold cross-validation accuracy ranged from 62.40% to 91.48%, with an average of 76.21%. The stacked classifier’s 5-fold cross-validation accuracy ranged from 84.95% to 91.00%, with an average of 88.60%. Lastly, the MLP using linear SVM features had a loss in the 5-fold cross-validation ranging from 0.2433 to 0.5151, with an average of 0.3355. The MLP using RBF SVM features had a loss in the 5-fold cross-validation ranging from 0.2221 to 0.6051, with an average of 0.3627. During the training, all classifiers showed a balance between precision and recall, and the LR classifier and the SVM with an RBF kernel had the optimal averaged precision (Figure. 1D).

The visualization of the classifier’s prediction and misclassification in training and testing sets showed that the misclassified cells mainly existed in two subsets of the CITE-seq data, Leiden clusters 1 and (C1, C3), where CD45RA^+^ and CD45RA^−^ cells were mixed by general clustering (Figure. 1B-C, 2A-B). The expression profile showed that the CCR7^+^SELL^+^IL7R^+^TCF7^+^ cluster 1 was likely to be naive T cells or TCMs, and the CCR7^−^SELL^−^IL7R^low^KLRG1^+^NKG7^+^ cluster 3 had an obvious TEM/TEMRA similarity (Figure. 2C-D)[13]. As discussed previously, these 2 pairs of cell types are difficult to classify in scRNA-seq analysis because of the overlap in the expression patterns and the absence of CD45RA, so it is reasonable that we observed misclassifications mainly came from these two clusters. Nevertheless, all 6 classifiers were still able to correctly label the majority of the hard-to-classify cells (Table 6).

**Table 6:**
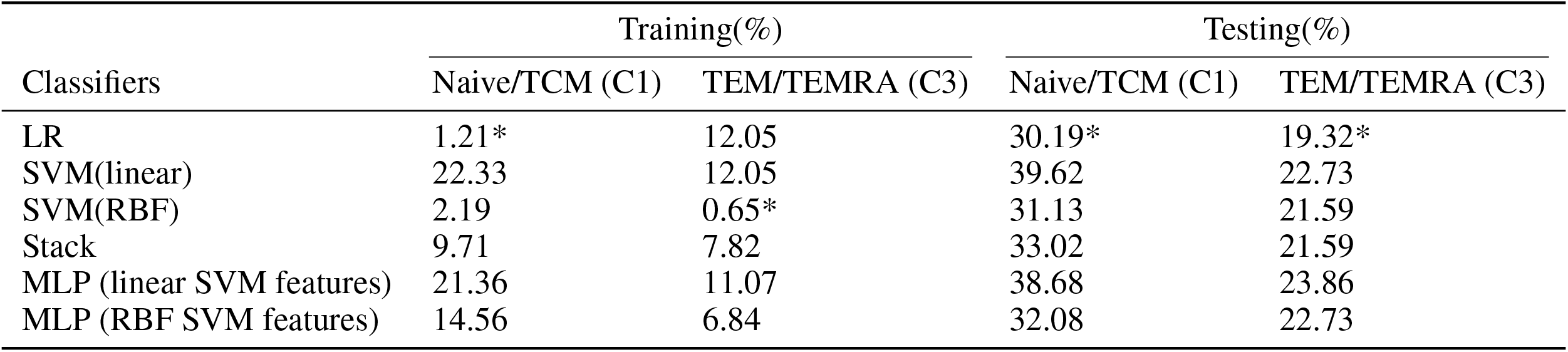
Percentage of Misclassification in Hard-to-Classify Clusters (Training&Testing)

| Classifiers | Training(%) |  | Testing(%) |  |
| --- | --- | --- | --- | --- |
|  | Naive/TCM (C1) | TEM/TEMRA (C3) | Naive/TCM (C1) | TEM/TEMRA (C3) |
| LR | 1.21* | 12.05 | 30.19* | 19.32* |
| SVM(linear) | 22.33 | 12.05 | 39.62 | 22.73 |
| SVM(RBF) | 2.19 | 0.65* | 31.13 | 21.59 |
| Stack | 9.71 | 7.82 | 33.02 | 21.59 |
| MLP (linear SVM features) | 21.36 | 11.07 | 38.68 | 23.86 |
| MLP (RBF SVM features) | 14.56 | 6.84 | 32.08 | 22.73 |

**Figure 2.**
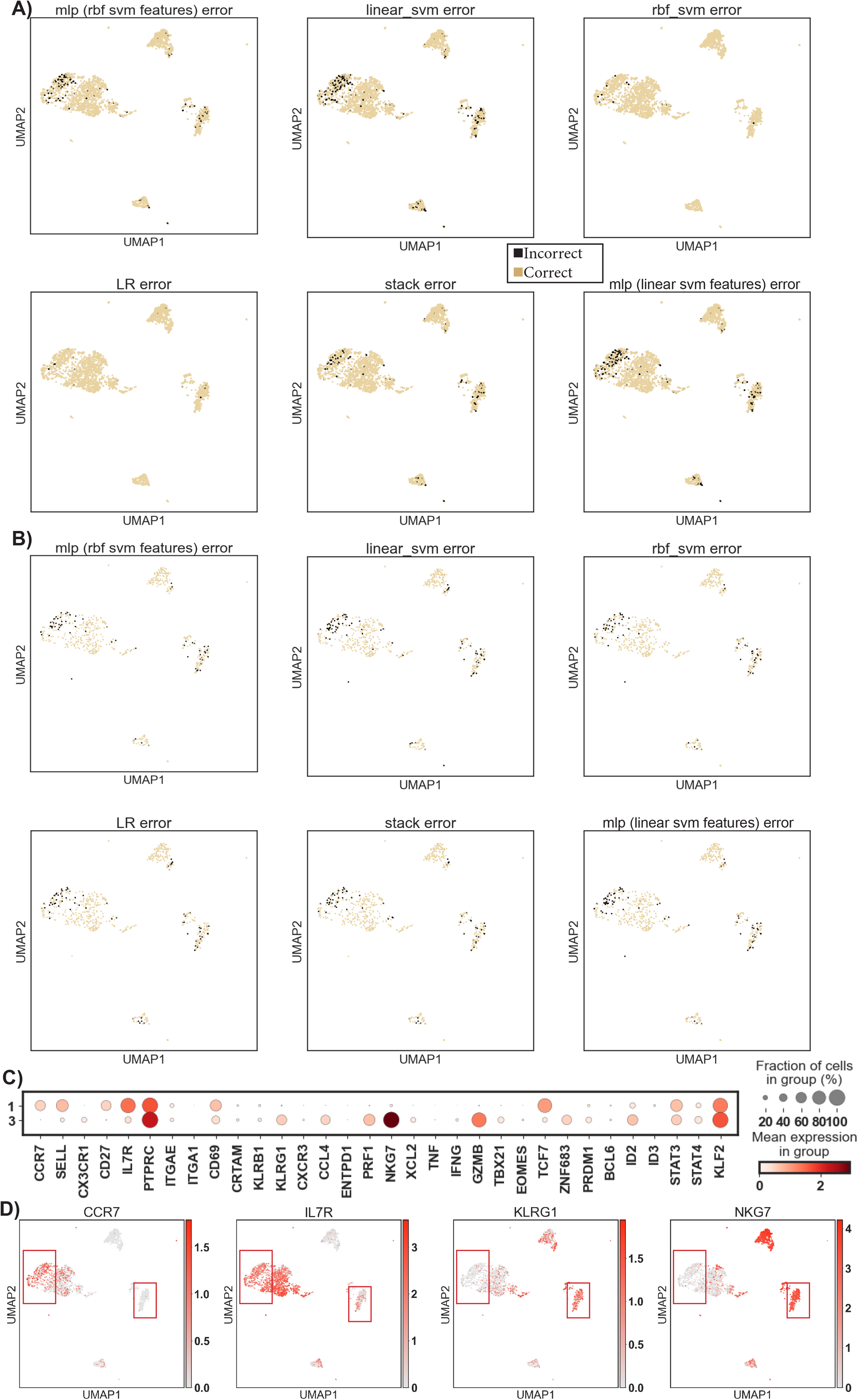
Misclassification and hard-to-classify clusters in the training/testing CITE-seq dataset. A), B) Visualization of classifiers’ wrong predictions (misclassification) in the A) training data, and B) testing data embedded on the UMAP coordinates. The first subplot in each plot shows the Leiden clustering as a reference. C) Expression (log-transformed corrected counts) of well-studied T marker genes in clusters 1 and 3 of the CITE-seq data. D) Visualization of T subsets marker expression in CITE-seq data on UMAP.

Applying the classifiers to a CD4^+^CD45RA^−^ unseen dataset allowed the evaluation of their performance when the input data is generated from a different experiment. Two SVMs and the MLP using linear SVM features achieved a high accuracy (>99%), and for the less accurate classifiers, LR and stacked models, the misclassifications were mainly from Leiden clusters 5, 10, and 11 (C5, C10, C11) (Figure. 3A, Table 3). Gene expression profiles of these 3 clusters showed they were likely to be resting (C5, C10; PRF1^−^GZMB^−^NKG7^−^) and activated (C11; PRF1^+^GZMB^+^NKG7^+^) CD4^+^ TEMs given their CCR7^−^SELL^−^IL7R^+^TNF^+^IFNG^+^ pattern (Figure. 3B-C). Indeed, CD4^+^ TEMs are also found to be able to re-express CD45RA[49], which makes these 3 clusters hard-to-classify if their CD45RA level is unknown. With that being said, the SVM with an RBF kernel perfectly labeled cells from these clusters based on their gene expression level (Table 7), and the other SVM with a linear kernel and the MLP using linear SVM features also performed well.

**Table 7:** Percentage of Misclassification in Hard-to-Classify Clusters (Unseen)

| Classifiers | Unseen(%) |  |  |
| --- | --- | --- | --- |
|  | TEM Act (C5) | TEM Rest 1 (C10) | TEM Rest 1 (C11) |
| LR | 47.57 | 27.13 | 79.81 |
| SVM(linear) | 3.80 | 0.39 | 2.12 |
| SVM(RBF) | 0* | 0* | 0* |
| Stack | 36.52 | 13.44 | 70.58 |
| MLP (linear SVM features) | 0.82 | 0.39 | 2.50 |
| MLP (RBF SVM features) | 13.93 | 2.97 | 47.5 |

**Figure 3.**
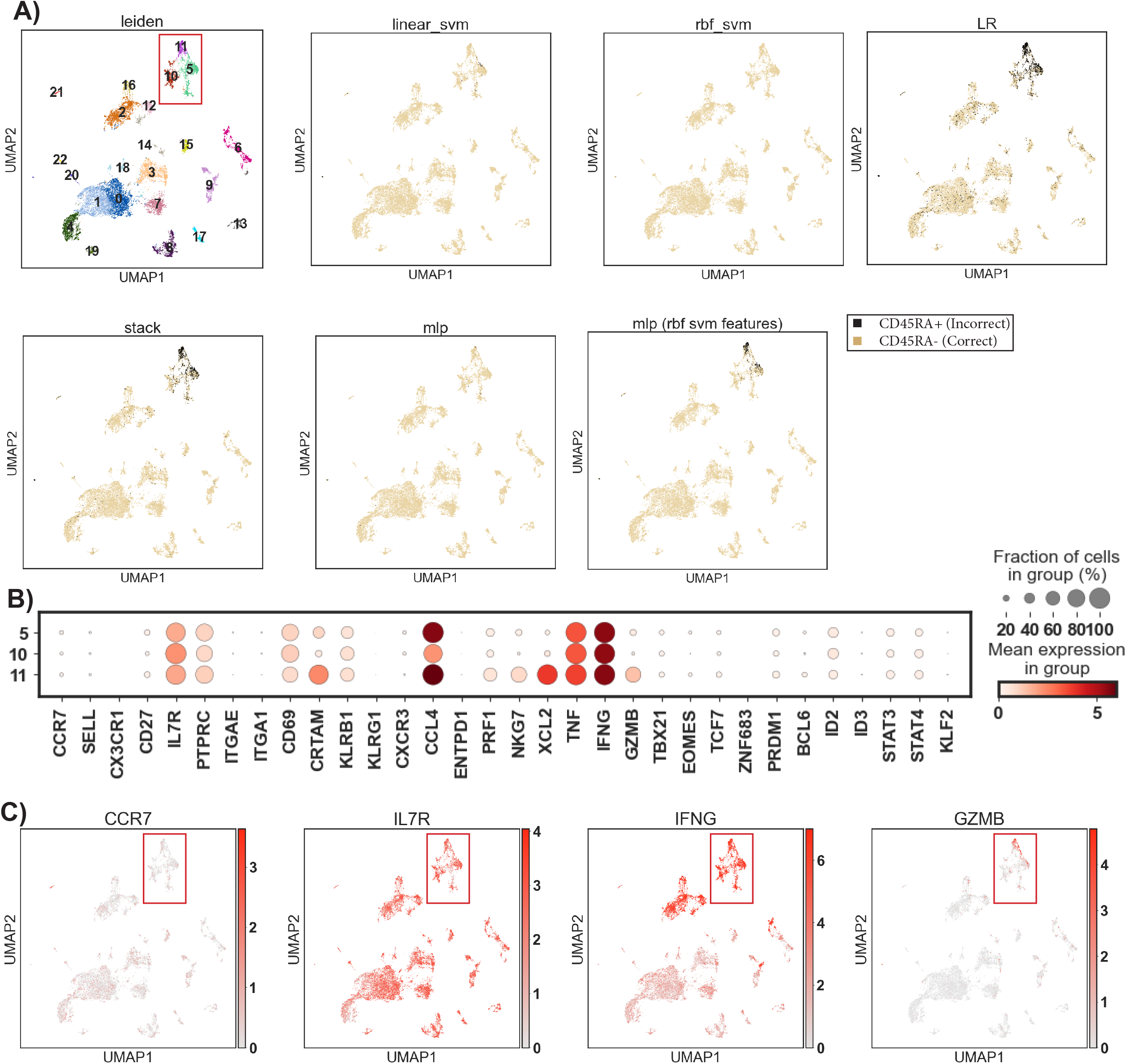
Misclassification and hard-to-classify clusters in the unseen dataset. A) Visualization of classifiers’ predictions in the unseen data embedded in the UMAP coordinates. Given the unseen data was reported as all CD45RA-, which should not have any CD45RA+ cells, this plot also visualizes classifiers’ wrong predictions (misclassification) in the unseen data. The first subplot in each plot shows the Leiden clustering as a reference. B) Expression (log-transformed corrected counts) of well-studied T marker genes in clusters 1 and 3 of the CITE-seq data. C) Visualization of T subsets marker expression in the unseen data on UMAP.

## 4 Discussion

This study explored several simple yet robust machine-learning classifiers to predict the CD45RA level of cells from its NGS scRNA-seq count matrix. We reported genes that were differentially expressed between CD45RA^+/-^, and we suggested that the transcription of genes like NPDC1 and AQP3 could be potentially indicative of the CD45RA protein level in human T cells. We trained and optimized several classifiers and compared their performance in making an accurate prediction. Although all of them had acceptable accuracy, two non-linear classifiers, SVM with RBF kernel and MLP with ReLu and Sigmoid activation functions (using linear SVM features), performed the best and almost perfectly predicted the CD45RA labels of unseen cells.

The key advantage of this classifier is its ability to overcome the technical limitations associated with short-read sequencing, which has historically struggled with detecting CD45RA and isoform identification. Consequently, our classifier addresses an unmet need in single-cell transcriptomics and provides a simple yet valuable tool for immunologists. Recent advances in long-read sequencing technologies, such as Oxford Nanopore, have shown promise in addressing some of the limitations of short-read sequencing, especially in resolving complex isoforms and alternative splicing events[50]. There is ongoing research and development to explore the use of long-read sequencing technologies, like Oxford Nanopore, for single-cell RNA-seq[51]. However, despite these advances, long-read sequencing remains relatively expensive and less accessible for many researchers. Isoform prediction from raw sequencing reads has been an area of active research, with methods such as StringTie[52] and Cufflinks[53] developed to address this challenge. While these methods have demonstrated success in predicting isoforms, they can be computationally intensive and may require specialized expertise.

The foundation of this study is the CITE-seq data, which simultaneously profiled the transcriptome and surface proteins at single-cell resolution [16]. As the technology matures and becomes more accessible, it is likely that the increased adoption of CITE-seq in the future will address the issue of CD45RA reporting as well as the identification of other difficult-to-report markers. However, since it is a relatively new technique that uses DNA-barcoded antibodies for protein detection, which increases the cost and complexity of the experiment, it has not yet become as popular as the conventional scRNA-seq. Therefore, it is anticipated that more conventional T-cell scRNA-seq experiments will still be conducted. In this way, our model can provide insights into analyzing existing and incoming conventional scRNA-seq datasets easily and cost-effectively.

In conclusion, our study presents a novel CD45RA+/-binary classifier that addresses the challenges associated with short-read sequencing and provides an efficient solution for immunologists working with scRNA-seq data. Future work could focus on refining the classifier’s performance and extending its applicability to other cell markers or transcriptomic technologies. Moreover, integration with existing bioinformatics pipelines and tools could enhance its utility and enable researchers to uncover novel insights into the complex world of immune cell biology.

## 5 Data Availability

The resultant package ScCD45RA can be found at https://github.com/WeldonSchool-BrubakerLab/ScCD45RA and can be installed from the Python Package Index (PyPI) using the command “pip install sccd45ra”.

## 6 Acknowledgements

The authors would like to thank Vivek Gupta (Purdue University, Department of Computer Science) for his support and advice.

## 7 Author contributions

Conceptualization: R.R.; Methodology: R.R., D.B.; Analysis: R.R.; Writing–original draft: R.R.; Writing–review & editing: R.R., D.B.; Administration: D.B.

